# Pattern Detection in Multiple Genome Sequences with Applications: The Case of All SARS-CoV-2 Complete Variants

**DOI:** 10.1101/2021.04.14.439840

**Authors:** Konstantinos F. Xylogiannopoulos

## Abstract

Pattern detection and string matching are fundamental problems in computer science and the accelerated expansion of bioinformatics and computational biology have made them a core topic for both disciplines. The SARS-CoV-2 pandemic has made such problems more demanding with hundreds or thousands of new genome variants discovered every week, because of constant mutations, and the need for fast and accurate analyses. Medicines and, mostly, vaccines must be altered to adapt and efficiently address mutations. The need of computational tools for genomic analysis, such as sequence alignment, is very important, although, in most cases the resources and computational power needed is vast. The presented data structures and algorithms, specifically built for text mining and pattern detection, can help to address efficiently several bioinformatics problems. With a single execution of advanced algorithms, with limited space and time complexity, it is possible to acquire knowledge on all repeated patterns that exist in multiple genome sequences and this information can be used for further meta analyses. The potentials of the presented solutions are demonstrated with the analysis of more than 55,000 SARS-CoV-2 genome sequences (collected on March 10, 2021) and the detection of all repeated patterns with length up to 60 nucleotides in these sequences, something practically impossible with other algorithms due to its complexity. These results can be used to help provide answers to questions such as all variants common patterns, sequence alignment, palindromes and tandem repeats detection, genome comparisons, etc.

## I. Introduction

The current COVID-19 pandemic has turned all the lights, commercial, scientific, political towards the biotechnology industry and its efforts to address as soon as possible the virus consequences. Major pharmaceutical companies worldwide have invested enormous amounts in new technologies for the past couple of decades and the first promising results from technologies such the mRNA vaccines have become visible. Indeed, the fast expansion of the biotechnology industry with the help of advanced computing infrastructures, such as cloud computing, has opened a new era in the domain.

Since the beginning of computer science some of the most common problems addressed were related to pattern matching and searching for bioinformatics. There is a plethora of completely diverse methodologies and algorithms since early 1970 that were developed to deal with the simplest problems, such as to determine if a specific string exists in a biological sequence, to more complex such as multiple sequence alignment. Furthermore, the development of artificial intelligence and deep learning provided more sophisticated tools for image analysis or clinical data analytics.

The analysis of biological sequences such as DNA, RNA, proteins, etc. it is considered a standard string problem in computer science since such sequences are built from predefined discrete alphabets like nucleotides or amino-acids encoding. What make these string problems to be challenging in bioinformatics and computational biology, from mathematical and computer science perspective, is the size of the strings and the computationally intensive procedures to answer them, which in some cases cannot provide solutions in short time and with regular resources. For example, the human genome, a 3.1GB long string, it was initially sequenced in 2001 [1] and it was practically impossible to be analyzed as a single piece of information since only supercomputers could keep on memory such long strings. Nowadays, advanced hardware and clustering framework systems are used for such analyses since, for example, the construction of a suffix tree, just for the first human chromosome with size 270MB, requires 26GB of memory [4]. New technologies, for instance, Next Generation Sequencing (NGS) from top leading companies require advanced computational tools and algorithms, specifically designed for string matching problems in order to perform sequence alignment in multiple (usually millions) genomic fragments simultaneously.

In [31] it was presented for first time the analysis of the full human genome with the detection of all repeated patterns. However, that initial attempt was just a proof of concept and technology. The current work will present that is possible, with limited resources and in short time, to analyze thousands of complete genomes and detect all repeated patterns that exist in them. Moreover, it will be presented how the combination of an advanced data structure and the results of such analysis can help to answer many pattern detection related problems. Finally, the possibilities and the potentials of the tools described to be used in specific type of string problems with applications on a large dataset comprised from all SARS-CoV-2 full genome variants recorded on March 10, 2021, will be presented.

In order to achieve such results, the Multivariate Longest Expected Repeated Pattern Reduced Suffix Array (LERP-RSA) data structure will be used in combination with the All Repeated Patterns Detection (ARPaD) algorithm [26], [27], [28]. In brief LERP-RSA is a variation of the standard Suffix Array [25] data structure using though the actual, lexicographically sorted, suffix strings. The ARPaD algorithm, both in its recursive and non-recursive variant, has the ability to scan only once the LERP-RSA and detect every pattern occurs at least twice in it. Additionally, the algorithm is pattern agnostic, which means that it requires no input rather it scans the data structure once and returns all results in a deterministic way regardless of string or pattern attributes, i.e., frequency, length, alphabet, overlapping or not, etc.

So far, LERP-RSA and ARPaD have been extensively used in many, diversified, domains with exceptional results, regardless of hardware limitations and the vast datasets in most cases, making them a state-of-the-art for big data problems in text mining and pattern detection [28]. As an example of resource demanding process in the bioinformatics domain, it is worth mentioning that an alignment between all complete SARS-CoV-2 genome variants was tried on the National Center for Biotechnology Information website, which holds the genome dataset, to receive a message that for more than 500 sequences alignment the user has to download the dataset and use own resources to perform the alignment.

The contribution of the current work is the analysis of more than 55,000 SARS-CoV-2 genome variants discovering all repeated patterns. These initial results have been used for further meta analytics, for example, discovering the longest pattern, with length 15 nucleotides, that exists among every variant of SARS-CoV-2, comparison among different organisms such as MERS, SARS and Human, identifying every frequent and infrequent pattern exists, etc. Additional applications are sequence alignment, detection of special attributes patterns such as palindromes and tandem repeats, etc. The proposed methods though, have also limitations such as the need for the sequence analyzed to have specific properties and in cases of simple, specific, problems this process could also be more time consuming. However, the benefit of using them on many diverse problems concurrently can overcome any initial hesitation.

The rest of the paper is organized as follows: Section II presents related work in string matching. Section III defines the problem and gives the motivation behind it. Section IV presents the proposed data structures and algorithms for pattern detection in biological sequences. Section V presents several applications conducted on the available dataset of all, complete SARS-CoV-2 variants and discusses the corresponding results. Finally, Section VI presents the conclusions and future extensions of the presented work.

### II. Related Work

From the very early stages of bioinformatics and the use of computers to perform biological sequence analyses, string matching problems had a crucial role. Many new algorithms and methodologies are presented every year that improve older approaches or introduce new [1], [3], [4]. Mainly, these methods and algorithms can be classified into two broad categories, the exact matching and the approximate matching [1], [4]. The first category is related to string problems where we seek to find patterns matching entirely the input string such as, for example, specific sequence matching a protein transcription promoter. The second category can be much more complicated since many mutations, insertions, deletions and base changes may have occur making exact matching difficult, yet, very important, for example, to detect codon sequences which can produce the same protein.

More precisely, exact matching algorithms have dominated the field since early ‘70s. Many different approaches have been developed such as character or index based. This kind of methodologies include brute force algorithms where characters of the matching pattern are directly compared to the reference sequence. This leads to heavy computational algorithms, mainly because of the absence of any preprocessing and special data structures. The standards for such algorithms are the Boyer-Moore algorithm, usually used as a benchmark for efficiency measurement, that uses a shifting step based on a table holding information about mismatch occurrences and the Knuth-Morris-Pratt algorithm that uses a supplementary table to record temporal information during execution [1], [3], [4], [5], [6]. Another algorithm, variation of the first one mentioned, is the Boyer-Moore-Smith [7] while another extension is the Apostolico-Giancarlo algorithm based on both of the BM and KMP algorithms [8]. Additionally, we have the Raita algorithm based on dependencies that occur among successive characters [9]. More recent algorithms are the BBQ algorithm which introduces parallel pointers that perform searching from opposite directions [10] and several hybrid methods such as the KMPBS [11] and Cao et al. using statistical inference [12].

Except the brute force algorithms we have another important category, the hashed based [1], [3], [4]. Such algorithms are based on the hashing concept in order to produce hashing values and compare patterns rather than performing a direct character comparison. The main benefit from such approach is the considerable improvement of calculation time [13], yet, as with most hashing algorithms, they suffer from the hashing collision problem. Typical examples of such algorithms is the Karp-Rabin which is based on modular arithmetic to perform hashing [14] and the Lecroq algorithm, which first splits the sequence to subsequences and then the pattern matching is performed on each sequence [15]. Classic algorithms are also the non q-gram algorithms such as the Wu and Manber [16] where the searching pattern is completely encoded for pattern matching purposes. Furthermore, more recently developed algorithms are the multi-window integer comparison algorithm based on suffix strings data structures such as the Franek-Jennings-Smyth string matching algorithm [18] and the automata skipping algorithm developed by Masaki et al. [17]. More advanced hybrid approaches have also been presented that combine best practices from different approaches in order to optimize their performance such as, for example, Navarro’s algorithm [19] which can bypass characters using suffix.

A very well known and heavily used algorithm is implemented and used by the National Center for Biotechnology Information (NCBI). The Basic Local Alignment Search Tool (BLAST) and its variants [21] is used for comparing basic sequences, such as nucleotides sequences, found in DNA and/or RNA. The algorithm takes as inputs the desired string to search and the sequence to search into. Additionally, BLAST can execute inexact string matching, something usually extremely computationally intensive, for multiple sequence alignment purposes. Another algorithm, more accurate than BLAST, yet, more resources hungry and slower, is the Smith-Waterman algorithm [20].

An important aspect of pattern detection is the discovery of specific type of patterns in biological sequences such as palindromes and tandem repeats. The importance of such discoveries can be presented with one of the latest marbles in biology, the discovery of the clustered regularly interspaced short palindromic repeats (CRISPR) in bacteria and the use of CRISPR-Cas9 protein that allows to interfere with DNA in a molecular level [22]. Another well studied problem is the detection of short tandem repeats, something very difficult over whole genome. This kind of repeats are classic examples of repeats in protein encoding regions and are closely related to serious diseases, such as the Huntington’s disease [23]. An example of methods for tandems detection can be found in [23] which is based on DNA alignment using LAST software.

## III. Problem Definition

So far, we have presented several algorithms that are used in bioinformatics and computational biology. Yet, all these algorithms have as a common attribute the input pattern that is under investigation. Such type of algorithms can address specific problems and require each time to access the full dataset of one or more sequences to operate and produce results, which is inefficient.

To address bioinformatics and computational biology problems, it would be more preferable to have a data structure or a database of information that can be used for as many as possible queries and be transformed to valuable knowledge. Moreover, the full process should be able to (a) be contacted on commodity computers with limited resources, (b) keep the cost low, (c) allow scale up to deal with larger datasets and (c) address several different problems concurrently.

## IV. Proposed Approaches

The approaches on biological problems, which they will be described in the next sections of the paper, are applications of the Longest Expected Repeated Pattern Reduced Suffix Array (LERP-RSA) data structure [26], [27], [28] and the related family of algorithms such as ARPaD, SPaD and MPaD that are specifically designed for the LERP-RSA [27], [28]. Several applications of the aforementioned data structure and algorithms will be presented, as a pipeline of execution, that can either extract useful information directly from the dataset or the results generated, or can be used as an input for other type of meta analytics in biological sequences.

### A. LERP-RSA Data Structure

The Longest Expected Repeated Pattern Reduced Suffix Array (LERP-RSA) is a special purpose data structure for text mining and pattern detection, which has been developed and optimized to work with a variety of algorithms, with applications in many domains. Manber and Myers [25] defined the suffix array of a string as the array of the indexes of the lexicographically sorted suffix strings, which allows to perform several tasks on the string, such as pattern matching. The LERP-RSA is a variation of the suffix array, yet, it uses the actual suffix string and not only the position indexes. Although this type of data structure can have quadratic space complexity, which was one of the first disadvantage to bypass, with the use of the LERP reduction the data structure space complexity can be optimized to log-linear with regard to the input string. This has been proved with the Probabilistic Existence of Longest Expected Repeated Pattern Theorem [27], [28] that can be briefly stated as follows:

#### Theorem

If a string is considerably long and random and a pattern is reasonably long then the probability that the pattern repeats in the string is extremely small.

The theorem builds us the necessary foundation to calculate the longest expected repeated pattern given a very small probability that a repeated pattern exists with longer length. Therefore, the length of the suffix strings used to create the LERP-RSA, can be reduced significantly by using the following, briefly stated, Lemma [27], [28]:

#### Lemma

An upper bound for the Longest Expected Repeated Pattern (LERP) length given a probability *P(X)* in a string of length *n* with the use of an alphabet *Σ* of size *m* is:

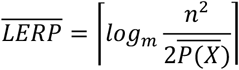

where *LERP* ≪ *n* and 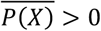.

The abovementioned Lemma is directly inducted from the Theorem and it has been proven in [27], [28]. Of course, the calculation of the longest repeated pattern can be performed by other methods, however, the use of the Lemma has some advantages since, e.g., building the suffix tree and determining the longest repeated pattern of a string on the suffix tree is a heavy computational process and in most of the cases it is impossible because of the string size. For example, in [28] the longest repeated patterns that exist in the first one trillion digits of π have been calculated, knowing in advance the upper limit for their length, while any other algorithmic approach is beyond any possibility with the currently available hardware. Yet, the Theorem and the Lemma have as a prerequisite that the string is random which limits the application for strings that do not have a random behavior. Briefly described, randomness means that every character of the alphabet occurs with the same frequency and this property should be valid for reasonably long substrings, following the normality of irrational numbers property as presented by the Calude’s Theorem [24]. Although this is true for most of the cases, unfortunately, biological sequences do not have random behavior and this problem can be solved with the MLERP process as it is described in [26], [28] and it has been used to analyze the full human genome [31].

The process of constructing the LERP-RSA with the use of the Lemma can be described with the following example. Let’s assume that the input string is *actactggtgt*. If we construct the array of the suffix strings then we will receive the structure of Fig.1.a where all suffix strings have been recorded, without sorting. Obviously, this structure has a quadratic space complexity of exact size *O(n(n* + 1) /2)or *O(n*^2^ ). If the size of the string becomes medium size, e.g., 10,000 characters, as an average human gene, then the space needed just to store the suffix strings, without sorting them, explodes to 100 million. What we can do to bypass the problem, for the initial example, is to reduce the size of the suffix strings to an arbitrary size to, e.g., five characters and create the structure of Fig.1.b. However, in this case we have the following to consider: (a) if the repeated patterns that exist and we want to discover are longer then we will miss all of them with length longer than the five characters and (b) if the repeated patterns are shorter then we are wasting space and time for sorting and analysis. This can be solved with the Lemma and the construction of the LERP-RSA of Fig.1.c, since if we reduce the size of suffix strings to three characters, for example, then the longest pattern that exists and is the *act* can be located at position 0 and 3. The use of value three is an example to illustrate the use the Lemma since it is not accurate for the specific, very sort, example. Since the LERP has length *O* (log *n*), with regard to the size of the input string, then the space complexity of the entire LERP-RSA is *O* (*n* log *n*).

**Fig.1.**
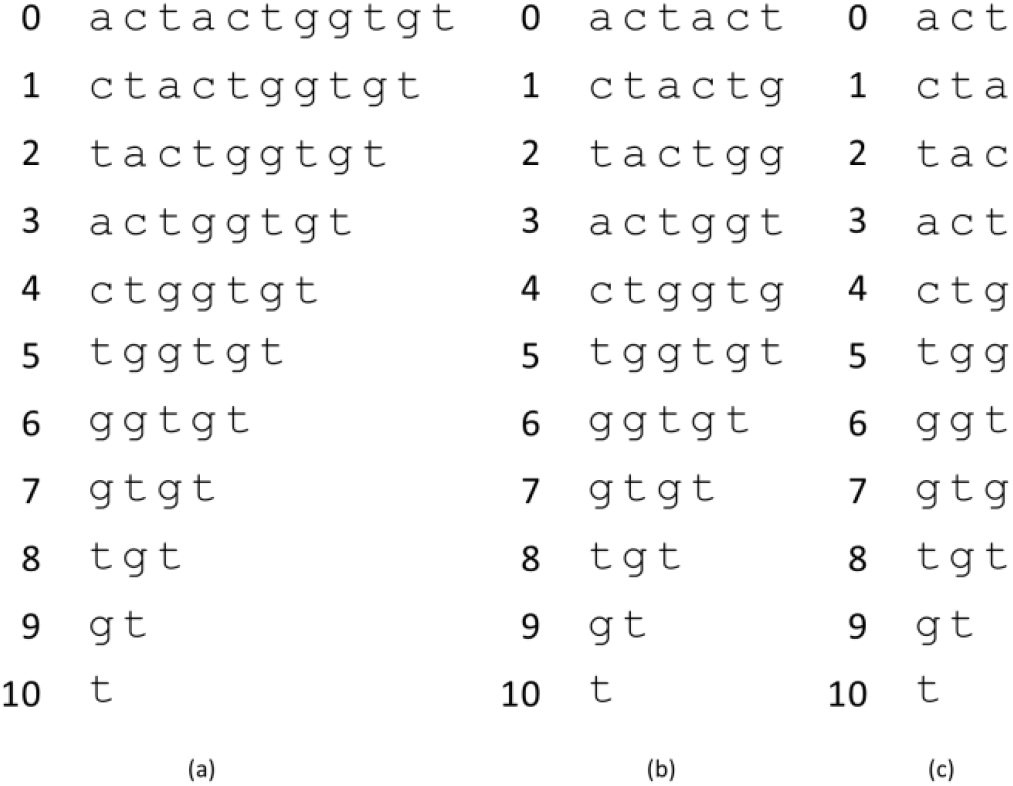
Suffix Array and Reduced Suffix Array for *actactggtgt*

The LERP-RSA data structure has some unique features that allows to be characterized as a state-of-the-art data structure for pattern detection and text mining purposes [27], [28]. These attributes are:

a. Classification based on the alphabet. The classification is determined by the Classification Level which is the power that the cardinality of the alphabet can be raised. For DNA sequences using the four nucleotides alphabet A, C, G and T, the classification can vary from one class, Σ^*CL*^ = 4^0^ = 1, for classification level zero to, e.g., 16 classes, Σ^*CL*^ = 4^2^ = 16, for classification level two with the construction of subclasses of suffix strings starting with *aa, ac, ag, at, ca, cc, cg, ct, ga, gc, gg, gt, ta, tc, tg* and *tt*. Therefore, instead of having one class we can have 16 with significantly smaller size each one, one sixteenth of the total if we assume equidistribution.
b. Network and cloud distribution based on the classes. Each class, regardless size, can be constructed or distributed independently over a local network or on the cloud. The classes can be stored and accessed when needed.
c. Full and semi parallelism. Since we have several, separate, classes the analysis and pattern detection algorithms can be executed on each class in parallel in full mode, all simultaneously, or semi-parallel mode where a block of classes is analyzed and when finished the analysis continues with the second block, etc.
d. Self-compression. When we use classification then we have in each class those suffix strings that specifically start with the class string. Therefore, the initial characters defining the class of the suffix strings in each class can be truncated and conserve space.
e. Indeterminacy. More space can be conserved for the cases that we do not care about the positions of the patterns rather than only for their existence. In this case the position indexes can be omitted.
f. The LERP-RSA can be constructed to describe multiple strings and allow the detection of patterns that exist not only in a single string but also among two or more different strings.

For many real world cases, such as biological sequences analysis and pattern detection, it is important to perform such tasks on multiple sequences. The last attribute described above is very important for these cases since it allows to detect patterns that are not repeated per se, yet, they exist once in several sequences, making them repeated. For this purpose, we need to construct the Multivariate LERP-RSA data structure as it can be described with the following example.

Let’s assume that we have two sequences *actactggtgt* and *ctactggtact* and, moreover, the LERP value is five while we have decided to use Classification Level two. In order to construct the data structure, we start with the first sequence at position zero and we use a sliding window of size five to determine the suffix strings (Fig.2.a). Additionally, for each position and suffix string we record the first two characters and we store the suffix string to the corresponding class (Fig.2.b). For example, the first five characters long substring of the first sequence is the *actac* and it will be stored in class *ac* with leading numbers to describe the sequence index (1, blue) and position in the specific sequence (0, black). We continue with the next substring *ctact* starting with *ct* which will be stored in class *ct*. We continue the process until position 9 where the substring *gt* with length two, exactly as the classification level, is the last one to be stored. The process of storing the suffix strings in each class (different colors for the example) can be performed directly or by sorting them. The same process repeats for the second sequence (Fig.3.a – Fig.3.b). Finally, the subclasses are combined together to create the lexicographically sorted Multivariate LERP-RSA (Fig.4.a) where each class is presented with different color.

**Fig.2.**
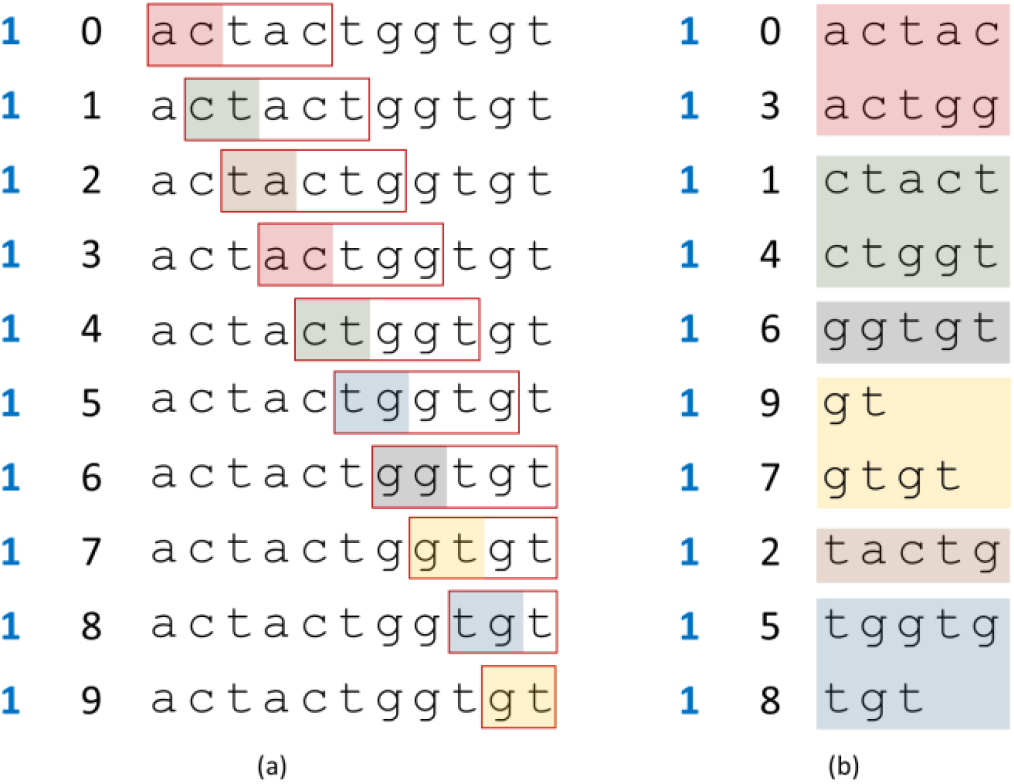
LERP-RSA construction for *actactggtgt*

**Fig.3.**
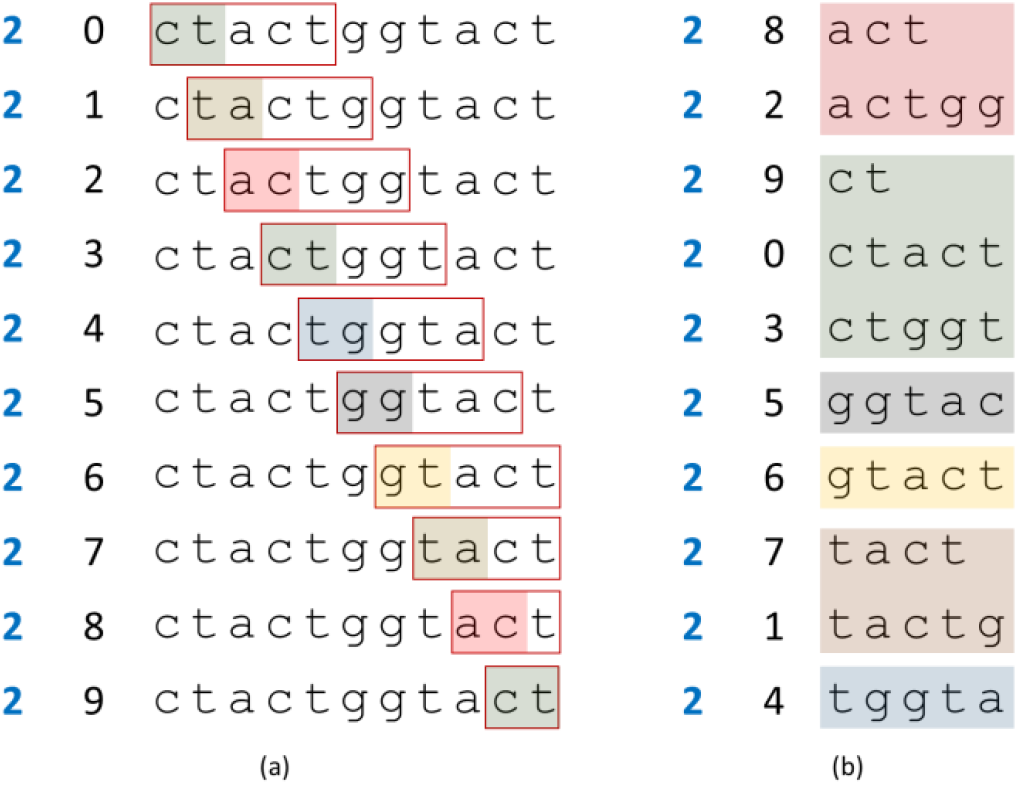
LERP-RSA construction for *ctactggtact*

**Fig.4.**
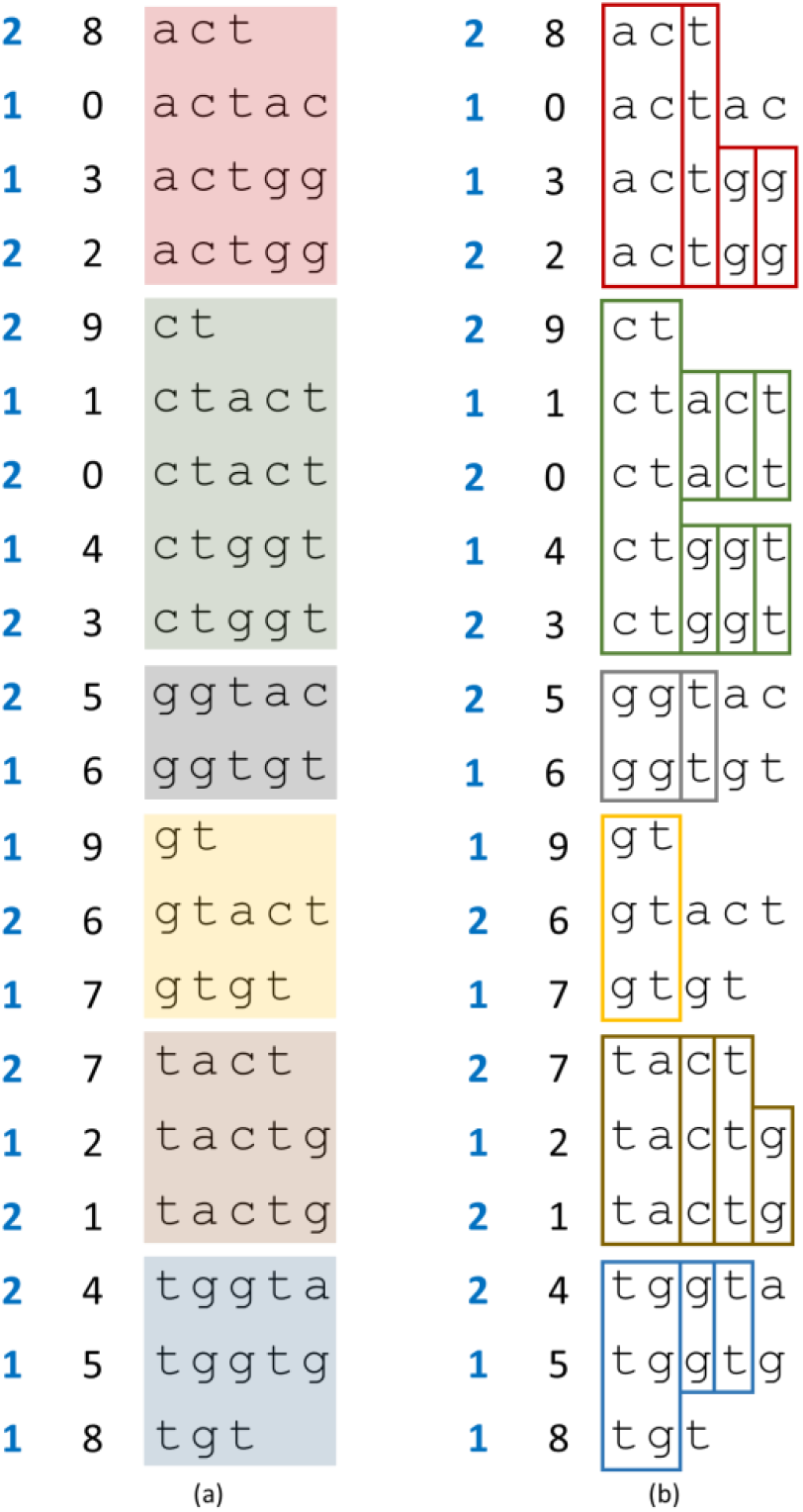
Multivariate LERP-RSA and ARPaD results

### B). ARPaD Algorithm

After constructing the Multivariate LERP-RSA data structure we execute the All Repeated Patterns Detection algorithm. The algorithm has two versions, the recursive left-to-right and the non-recursive top-to-bottom [28]. Both versions have the same time complexity *O*(*nlogn*). Since it is easier to present with an example the recursive, we will use the LERP-RSA of the previous subsection example in Fig.4.a.

First, the algorithm starts with the first class it has been created, *ac*, and counts how many strings starts with it (Fig. 4.b). Since there are four suffix strings in this class then then the class itself is a repeated pattern. The algorithm constructs a longer pattern with the first letter of the nucleotides alphabet, *a*, the *aca*. This does not exist and the algorithm continues with the other letters of the alphabet until it finds the pattern *act* which also appears four times (Fig.4.b). The process is repeated for longer patterns, starting with *acta*, until it finds the *actg* occurring twice and the longer *actgg* (Fig.4.b) which also occurs twice. With this the algorithm has discovered all repeated patterns of class *ac* or similarly starting with *ac*. The process is executed for each class and the ARPaD algorithm discovers at the end all repeated patterns (Fig.4.b). The non-recursive top-to-bottom version works in a similar way by comparing directly suffix string tuples.

Based on the above presented example, we can observe that ARPaD is executed on each class independently and, therefore, it can be executed in parallel. The only constrains for such execution is the available hardware, processors or cores and memory. For example, if the available resources do not allow for full parallel execution, we can start with the classes *ac* and *ta* which have the same number of suffix strings. Then we observe that class *ct* has five suffix strings while classes *gg* and *gt* have also five suffix strings combined. Therefore, we can execute in semi-parallel mode class *ct* with *gg* and when *gg* finishes, obviously before *ct*, we continue with class *gt*. This order of execution optimizes resources usage and minimizes idle time for the CPU.

Of course, we can execute ARPaD independently on each class, assuming enough resources. This can be achieved also for datasets that significantly exceed the available local resources by using the network and/or cloud distribution. This property of LERP-RSA and ARPaD allows to use completely isolated and diversified hardware, e.g., smartphones, to analyze each class in complete isolation from other classes instead of using expensive hardware infrastructure or clustering frameworks such as Hadoop and Spark.

### C. SPaD Algorithm

Another important algorithm of the ARPaD family is the Single Pattern Detection (SPaD) algorithm [28]. The SPaD algorithm is mainly used for meta-analyses purposes, when we want to discover specific information in the ARPaD results or LERP-RSA, and its correctness has been proven in [28]. Moreover, especially with the LERP-RSA it can be extremely efficient with time complexity *O(1)*with regard to the input string. Although ARPaD can be executed once to detect all repeated patterns that can be stored for later meta-analyses purposes, SPaD has to be used every time we need to, e.g., check the existence of non-repeated patterns. For this purpose, we execute the SPaD directly on the LERP-RSA data structure since single occurred patterns can exist only in the LERP-RSA, if they do exist. There are two distinct cases of SPaD execution with regard to the length of the pattern we need to find; if a pattern is equal or shorter than LERP or if a pattern is longer than LERP.

Using the previously stated example we can describe the SPaD algorithm using two sample patterns with regard to their size in comparison to the LERP value. The first pattern is the *gtg*, which is not repeated pattern since we cannot find it in the ARPaD results and it is shorter than the LERP value. Since the pattern starts with *gt*, SPaD starts in the appropriate *gt* class and using the binary search algorithm approach finds the suffix string in the class, *gtact* (Fig.5.a-1). Since *gtg* is lexicographically after *gtact*, the algorithm continues in the second half of the *gt* class and finds once the pattern in the suffix string *gtgt* (Fig.5.a-2). Therefore, the pattern *gtg* exists once in the first sequence at position seven.

**Fig.5.**
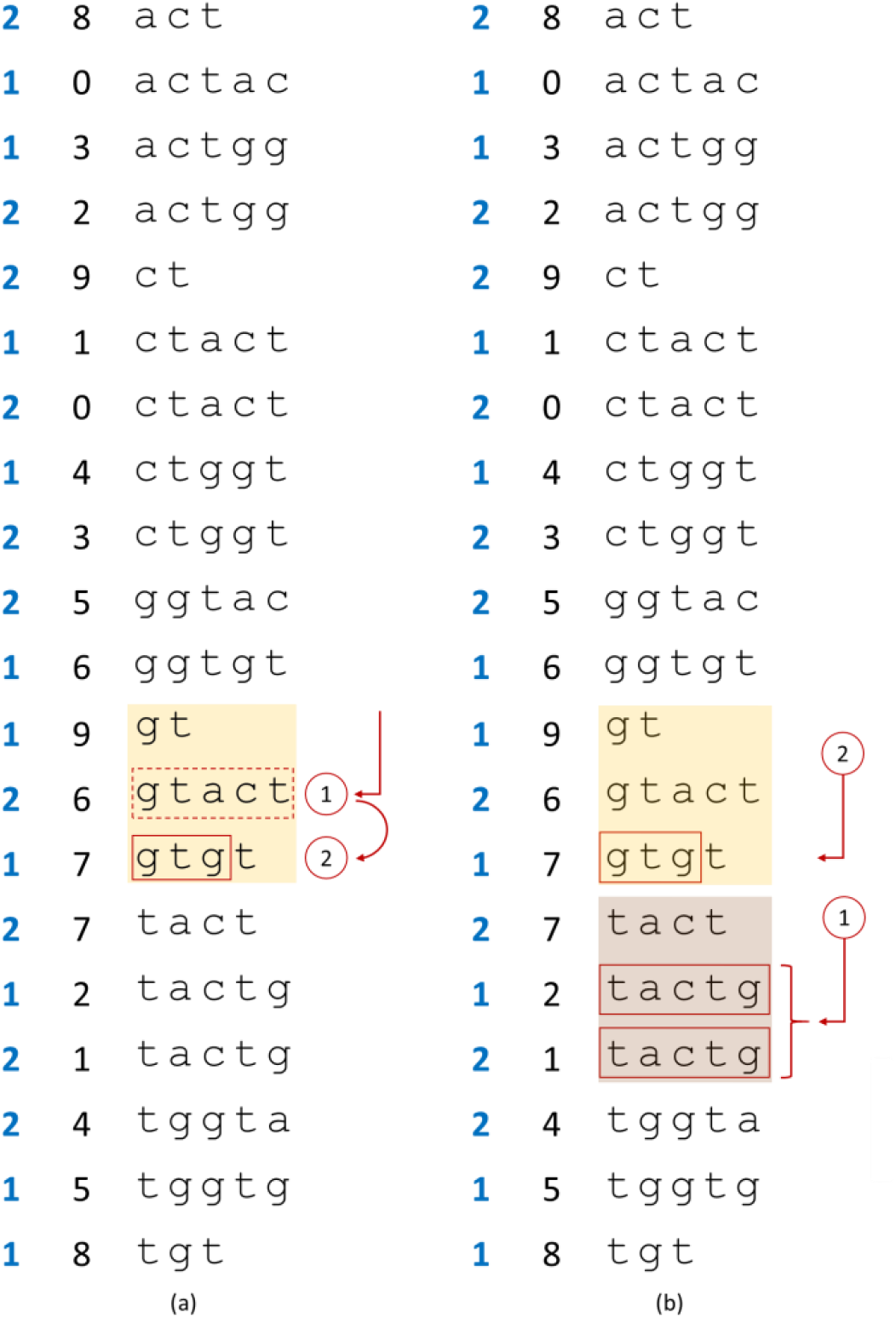
SPaD algorithm example for pattern *gtg* and *tactggtg*

The next example is the *tactggtg* pattern which is longer than the LERP value. The first step is to break down the pattern under investigation to fragments of size LERP, except, of course, the last one which can be smaller. Therefore, for the particular pattern we have two fragments, the *tactg* and the *gtg*. The next step for the SPaD algorithm is to search for each fragment and record if it exists and where (Fig.5.b). If at least one of the fragments do not exist in the LERP-RSA then, obviously, the pattern does not exist in any sequence. However, if we find all fragments to occur somewhere in the LERP-RSA then SPaD has to check if the full pattern exists. In order to perform this SPaD uses the Crossed Minimax Criterion [22]. For the specific example, we can observe that the first fragment exists twice in the class *tc* and, specifically, for the first sequence at position two and for the second sequence at position one (Fig.5.b-1). The second fragment can be found only once in class *gt* for sequence one at position seven (Fig.5.b-2). First of all, since the second fragment does not exist in the second sequence, therefore, the pattern does not exist in the second sequence. For the first sequence, the first fragment exists at position two and the second at position seven. Since the second position (7) is equal to the first position (2) plus LERP value (5), therefore, the pattern exists in the first sequence at position two.

The SPaD algorithm except of its straight forward application described above can also be used with wildcards or regular expressions, for the detection of more complex patterns. Let’s assume that we want to detect all patterns with the form *t??tg*, where the symbol *?* means any character from the alphabet. Therefore, we care to find patterns such as *t**aa**tg, t**ac**tg, t**ag**tg, t**at**gt, t**ca**gt*, etc. Executing the SPaD for each combination or by using regular expressions we can detect the patterns *tactg* at positions (1, 2) and (2, 1) and *tggtg* at position (1, 5). However, when we use wildcards or we need to detect more multiple patterns, the best option for optimization purposes, is the use of the MPaD algorithm of the next subsection.

### D. MPaD Algorithm

The Multiple Pattern Detection (MPaD) [28] algorithm is a direct extension of the SPaD. In the case of multiple pattern detection instead of using one time after the other the SPaD algorithm the process is optimized with the use of the MPaD. Practically, the first step of the SPaD is extended by breaking down all patterns into fragments and adding common fragments into batches. This can help the algorithm execution because patterns can have shared fragments that they will be searched only once and if not existed a complete batch of patterns can be rejected simultaneously, instead of repeating the process. As with SPaD, MPaD can also be used with wildcards for more advanced pattern detection.

### E. Metadata Analytics

After the completion of the data analysis several metadata analyses can be performed. These analyses depend on several factors and the problems that we want to address such as sequence alignment, genome comparison, palindromes and tandem repeats detection, etc. The importance of the full analysis and repeated patterns detection is that it needs to be executed only once and our further, detailed, meta analyses in the results are standalone processes. Moreover, the results can be stored on external storage media, locally or remotely on the cloud, and accessed whenever is needed, by class, without the need to repeat the analysis or access the full dataset.

### F. Synopsis

The first step of applying any of the proposed algorithms is the construction of the Multivariate LERP-RSA data structure. The LERP-RSA data structure construction has a space and time complexity of *O*(*n* log *n*) as it has been already discussed thoroughly. In the case of the Multivariate LERP-RSA, since we have *m* sequences of approximate length *n*, the total space complexity is *O*(*mn* log *n*)since the total size of the dataset, if it is considered a single sequence is *m × n*. However, the logarithmic part of the complexity is not equally *m × n* since the sequences are independent and according to Calude’s theorem [24] we do not expect such long repeated patterns.

When LERP-RSA construction is completed then we execute the All Repeated Patterns Detection (ARPaD) algorithm which is the second step of the methodology for data analytics and pattern detection in biological sequences. It is important to mention that both steps are executed once during the lifecycle of the data analytics process. ARPaD has time complexity *O*(*mn* log *n*) and the results can be stored for any kind of meta-analytics.

Having the LERP-RSA data structure and ARPaD results stored then we can use SPaD, MPaD or any other algorithm on the precalculated results to perform any kind of analysis such as sequence alignment, genomic comparisons, detecting primers for polymerase chain reaction process, identifying protein promoters, palindromes and tandem repeats, etc. The full process can be depicted with Fig.6.

**Fig.6.**
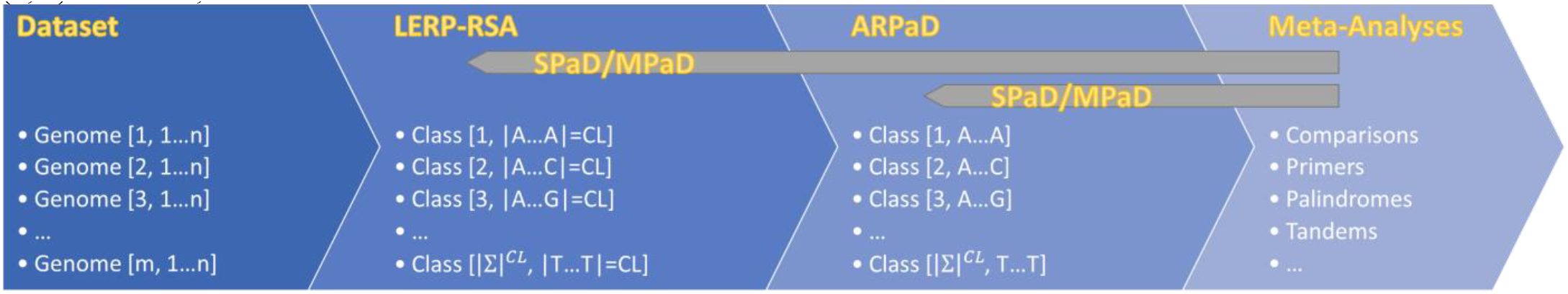
LERP-RSA, ARPaD, SPaD and MPaD process execution

## V. Experimental Analyses and Applications

For the presentation of possible applications of LERP-RSA and the ARPaD family algorithms on different use cases a dataset consisted from all SARS-CoV-2 complete genome variants has been used. The dataset was recorded on March 10^th^, 2021, and downloaded from the National Library of Medicine at the National Center for Biotechnology Information (NCBI) [30] in its FASTA format (taxid 2697049). The recorded dataset at the specific date consists of 55,733 sequences with an average sequence length of 29,812. However, there is one sequence, the MT873050.1/USA/MA-MGH-01491/2020, which has length just 2,859 bases and it has been removed from the dataset. The total size of the dataset is approximately 1.7GB, half the size of the total human genome.

Although SARS-CoV-2 is a single stranded RNA plus virus, the DNA reverse transcribed sequences have been recorded in the dataset. For this reason, the standard nucleotides alphabet {A, C, G, T} has been used and the sequence strings have been cleaned from many non standard characters such as N, R, W etc. and replaced with a neutral symbol $ to help avoid meaningless patterns.

For the analysis a laptop computer with an Intel i7 CPU at 2.6 GHz has been used with 16 GB RAM and an external disk of 1 TB for a semi-parallel execution, consuming approximately 7 hours. For a wider semi parallel execution, four computers with approximately same configuration have been used in order to execute per computer one master class of the alphabet (A$$, C$$, G$$ and T$$) and took approximately 2 hours. The Classification Level used is three, creating the 64 codon elements used for the translation process to proteins (AAA, AC, AAG, …, TTG, TTT). The results of this analysis are enormous and for practical reasons only few, interesting, use cases and metadata analyses will be presented here. The LERP value used is 60; 20 codons length. The total size of the LERP-RSA data structure on disk is 113GB, which practically means that it cannot be processed as a single class dataset. The larger class though, using the predefined classification, is the TTT with size approximately 4GB while the smallest is the CCG with size approximately 300MB.

A summary of the ARPaD results can be found on Table There are 64 patterns with length three, as many as the classes, yet, with length four there are 320 instead of the expected 256. This happens because of the patterns which include the characters replaced with the neutral symbol $ and practically alters the alphabet size to five characters. The cumulative number of patterns with length up to 60 characters is 36.2 million approximately and the total cumulative occurrences of these patterns is approximately 96.2 billion (Table I).

**TABLE I.** ARPaD Results Pattern and Occurrences Statistics

| L. | Patterns | Total Occurrences | Cumulative Patterns | Cumulative Occurrences |
| --- | --- | --- | --- | --- |
| 3 | 64 | 1,660,414,227 | 64 | 1,660,414,227 |
| 4 | 320 | 1,660,359,327 | 384 | 3,320,773,554 |
| 5 | 1,600 | 1,660,304,426 | 1,984 | 4,981,077,980 |
| 6 | 7,569 | 1,660,249,245 | 9,553 | 6,641,327,225 |
| 7 | 27,438 | 1,660,190,602 | 36,991 | 8,301,517,827 |
| 8 | 70,814 | 1,660,119,119 | 107,805 | 9,961,636,946 |
| 9 | 133,352 | 1,660,027,964 | 241,157 | 11,621,664,910 |
| 10 | 188,260 | 1,659,926,448 | 429,417 | 13,281,591,358 |
| 11 | 224,922 | 1,659,832,352 | 654,339 | 14,941,423,710 |
| 12 | 251,019 | 1,659,748,345 | 905,358 | 16,601,172,055 |
| 13 | 273,082 | 1,659,669,551 | 1,178,440 | 18,260,841,606 |
| 14 | 293,766 | 1,659,592,547 | 1,472,206 | 19,920,434,153 |
| 15 | 314,004 | 1,659,516,221 | 1,786,210 | 21,579,950,374 |
| 16 | 334,066 | 1,659,439,977 | 2,120,276 | 23,239,390,351 |
| 17 | 354,032 | 1,659,363,717 | 2,474,308 | 24,898,754,068 |
| 18 | 373,900 | 1,659,287,879 | 2,848,208 | 26,558,041,947 |
| 19 | 393,726 | 1,659,212,566 | 3,241,934 | 28,217,254,513 |
| 20 | 413,518 | 1,659,137,358 | 3,655,452 | 29,876,391,871 |
| 21 | 433,277 | 1,659,062,077 | 4,088,729 | 31,535,453,948 |
| 22 | 453,004 | 1,658,986,770 | 4,541,733 | 33,194,440,718 |
| 23 | 472,682 | 1,658,911,339 | 5,014,415 | 34,853,352,057 |
| 24 | 492,363 | 1,658,835,779 | 5,506,778 | 36,512,187,836 |
| 25 | 512,003 | 1,658,760,092 | 6,018,781 | 38,170,947,928 |
| 26 | 531,620 | 1,658,684,241 | 6,550,401 | 39,829,632,169 |
| 27 | 551,236 | 1,658,608,193 | 7,101,637 | 41,488,240,362 |
| 28 | 570,837 | 1,658,531,896 | 7,672,474 | 43,146,772,258 |
| 29 | 590,389 | 1,658,455,338 | 8,262,863 | 44,805,227,596 |
| 30 | 609,936 | 1,658,378,567 | 8,872,799 | 46,463,606,163 |
| 31 | 629,469 | 1,658,301,541 | 9,502,268 | 48,121,907,704 |
| 32 | 648,987 | 1,658,224,306 | 10,151,255 | 49,780,132,010 |
| 33 | 668,493 | 1,658,146,885 | 10,819,748 | 51,438,278,895 |
| 34 | 687,992 | 1,658,069,281 | 11,507,740 | 53,096,348,176 |
| 35 | 707,492 | 1,657,992,526 | 12,215,232 | 54,754,340,702 |
| 36 | 726,957 | 1,657,915,839 | 12,942,189 | 56,412,256,541 |
| 37 | 746,422 | 1,657,839,970 | 13,688,611 | 58,070,096,511 |
| 38 | 765,901 | 1,657,764,229 | 14,454,512 | 59,727,860,740 |
| 39 | 785,343 | 1,657,688,359 | 15,239,855 | 61,385,549,099 |
| 40 | 804,752 | 1,657,612,290 | 16,044,607 | 63,043,161,389 |
| 41 | 824,126 | 1,657,535,758 | 16,868,733 | 64,700,697,147 |
| 42 | 843,506 | 1,657,459,126 | 17,712,239 | 66,358,156,273 |
| 43 | 862,863 | 1,657,382,229 | 18,575,102 | 68,015,538,502 |
| 44 | 882,214 | 1,657,305,261 | 19,457,316 | 69,672,843,763 |
| 45 | 901,530 | 1,657,228,003 | 20,358,846 | 71,330,071,766 |
| 46 | 920,835 | 1,657,150,209 | 21,279,681 | 72,987,221,975 |
| 47 | 940,133 | 1,657,072,148 | 22,219,814 | 74,644,294,123 |
| 48 | 959,415 | 1,656,993,902 | 23,179,229 | 76,301,288,025 |
| 49 | 978,666 | 1,656,915,484 | 24,157,895 | 77,958,203,509 |
| 50 | 997,897 | 1,656,836,870 | 25,155,792 | 79,615,040,379 |
| 51 | 1,017,104 | 1,656,758,090 | 26,172,896 | 81,271,798,469 |
| 52 | 1,036,296 | 1,656,679,189 | 27,209,192 | 82,928,477,658 |
| 53 | 1,055,505 | 1,656,600,251 | 28,264,697 | 84,585,077,909 |
| 54 | 1,074,712 | 1,656,521,257 | 29,339,409 | 86,241,599,166 |
| 55 | 1,093,911 | 1,656,442,200 | 30,433,320 | 87,898,041,366 |
| 56 | 1,113,080 | 1,656,363,054 | 31,546,400 | 89,554,404,420 |
| 57 | 1,132,244 | 1,656,283,873 | 32,678,644 | 91,210,688,293 |
| 58 | 1,151,426 | 1,656,204,674 | 33,830,070 | 92,866,892,967 |
| 59 | 1,170,610 | 1,656,125,433 | 35,000,680 | 94,523,018,400 |
| 60 | 1,189,792 | 1,656,046,122 | 36,190,472 | 96,179,064,522 |

Table II presents the most frequent 60 characters long patterns from each one of the 64 classes. The patterns in the table are sorted based on the average positioning in all sequences (variants).

**TABLE II.**
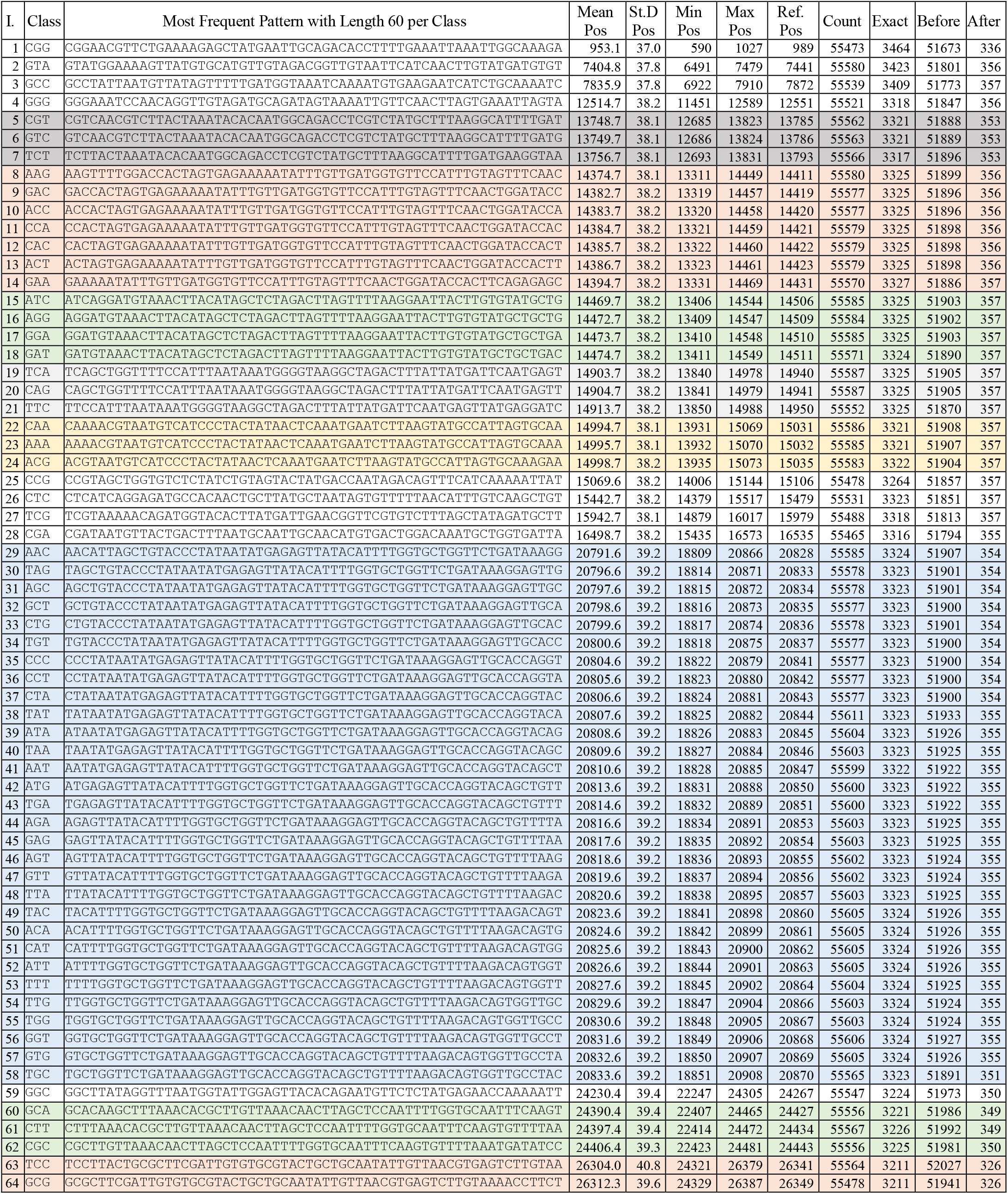
Positional descriptive statistics for most frequent patterns per class with length 60

The column next to mean positioning is the standard deviation of the pattern among all sequences, which takes values between 37 and 40 for all patterns. The next two columns are the minimum and maximum positions that the patterns have been detected in the sequences. The next column is the position that each pattern occurs in the reference sequence NC_045512.2. As we can observe, we can have some very interesting qualitative and quantitative information.

For example, for the first pattern in Table II for class CGG, we have in total 55,473 occurrences where 3,464 happen exactly at the same position as in the reference sequence while 51,673 happen before and 336 after. This can help us conclude that up to the specific position most of the variants (51,673) have more deletions than insertions in the genome while the rest (336) have more insertions than deletions.

In the same Table II, some patterns are marked with the same color. These variants are practically overlapping, as we can observe from their mean position which increments by one or a few more characters. These patterns can be further expanded with the use of other 60 characters long patterns or shorter patterns to form common regions in the sequences where most of the sequences are identical. Moreover, this information can be used for sequence alignment purposes, although it is a more demanding task, which will be presented in future work. The patterns in Table II create 17 different blocks in the SARS-CoV-2 sequence. These blocks practically separate the vast majority of the sequences to common regions and more blocks can be used with shorter patterns. It needs to be mentioned that this is not valid for all sequences since some may not occur in specific sequences due to mutations. Still, shorter patterns can reveal these blocks.

Another application of the proposed methodology is the comparison of genomes among different organisms. For example, in Table III we have all patterns from SARS-CoV-2 that exist at least once in every variant of the virus and has length greater or equal to 12. These patterns are compared with other organisms’ genomes such as the MERS virus (610 total variants, taxid 1335626), SARS virus (74,121 variants, taxid 694009) [30] and the human genome (GRCH38.p12) [29]. As we can observe, MERS virus has only one common pattern with SARS-CoV-2 that occurs in most of its variants. Yet, there are three more patterns that exist in one or two variants only, while all patterns exist practically in all variants of SARS virus, which it can be explained since SARS and SARS-CoV-2 belong to the same family of viruses. What it looks impressive is that all patterns exist in the human genome too, with different number of occurrences varying from 8 up to 930 but slightly longer patterns cannot be found in human genome. A possible application of this information is the determination of primers for PCR analyses. Since the patterns exist in all SARS-CoV-2 variants they can be used in pairs to amplify, practically, the largest part of the virus. However, if used with human DNA sample then PCR is not possible since human genome could also be amplified. This can be bypassed for the specific purpose with the use of longer patterns, e.g., with length 60 as the pattern in Table II, that do not exist in the human genome. Yet, since these patterns are not present in all SARS-CoV-2 variants two couples must be used that cover all possible cases. This can help to use PCR not just on specific SARS-CoV-2 proteins but on much larger parts of the genome. For example, if we use the 30 characters long patterns GTGCTGGTAGTACATTTATTAGTGATGAAG and GCGTGTAGCAGGTGACTCAGGTTTTGCTGC occurring at positions 934 and 27039 respectively, it is possible to amplify approximately 90% of the genome.

**TABLE III.**
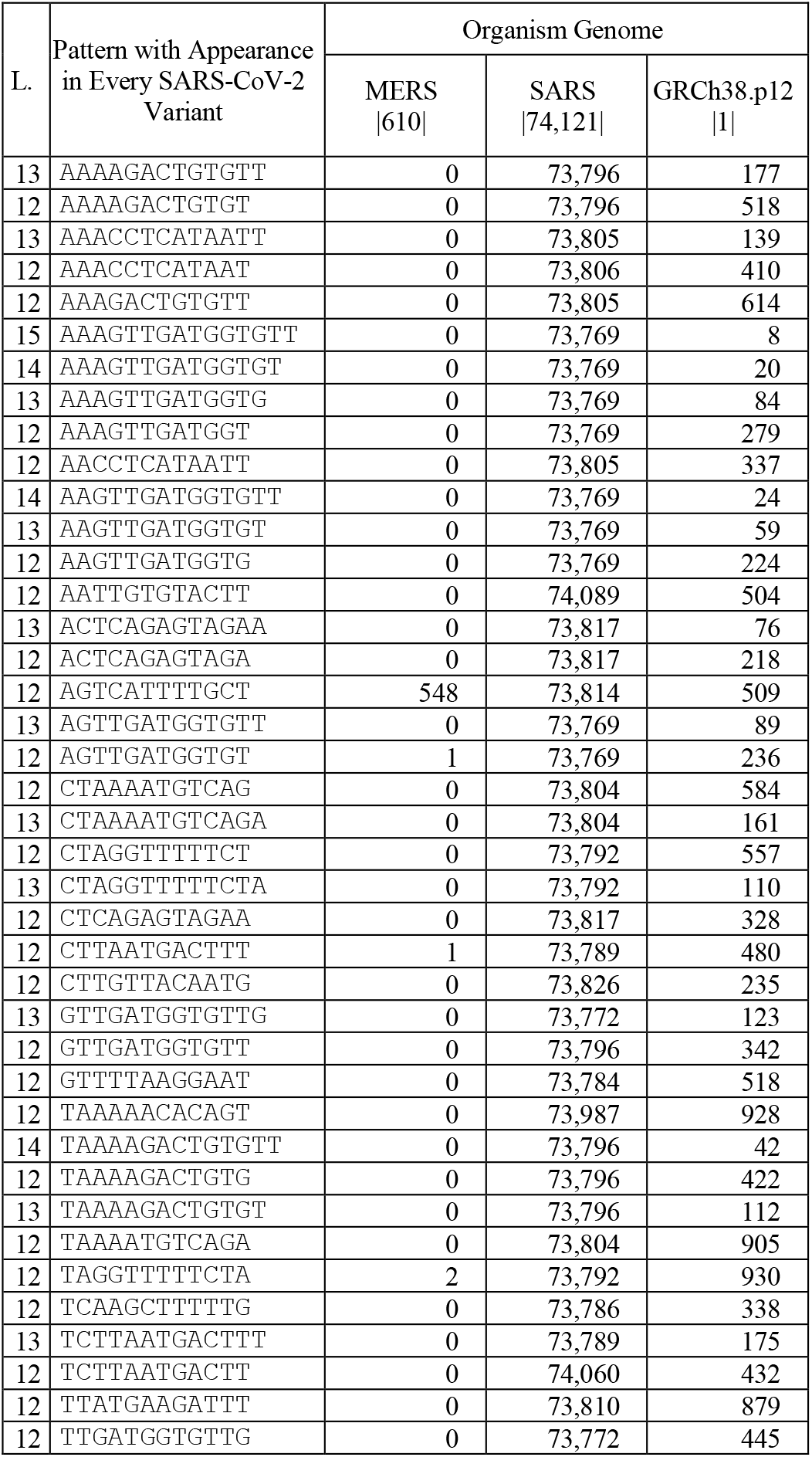
Comparison between different organisms

Finally, in Table IV some examples from palindromes and tandem repeats are presented. All the example patterns have been identified as repeated patterns and it is very easy to be filtered from the ARPaD results. The first six patterns present tandem repeats of total length eight or nine characters, constructed from tandems of length two, three or four characters. The next six patterns are palindromes of length eight and ten characters. Additionally, the occurring positions in the sequences are presented, truncated for patterns with many occurrences.

**TABLE IV.**
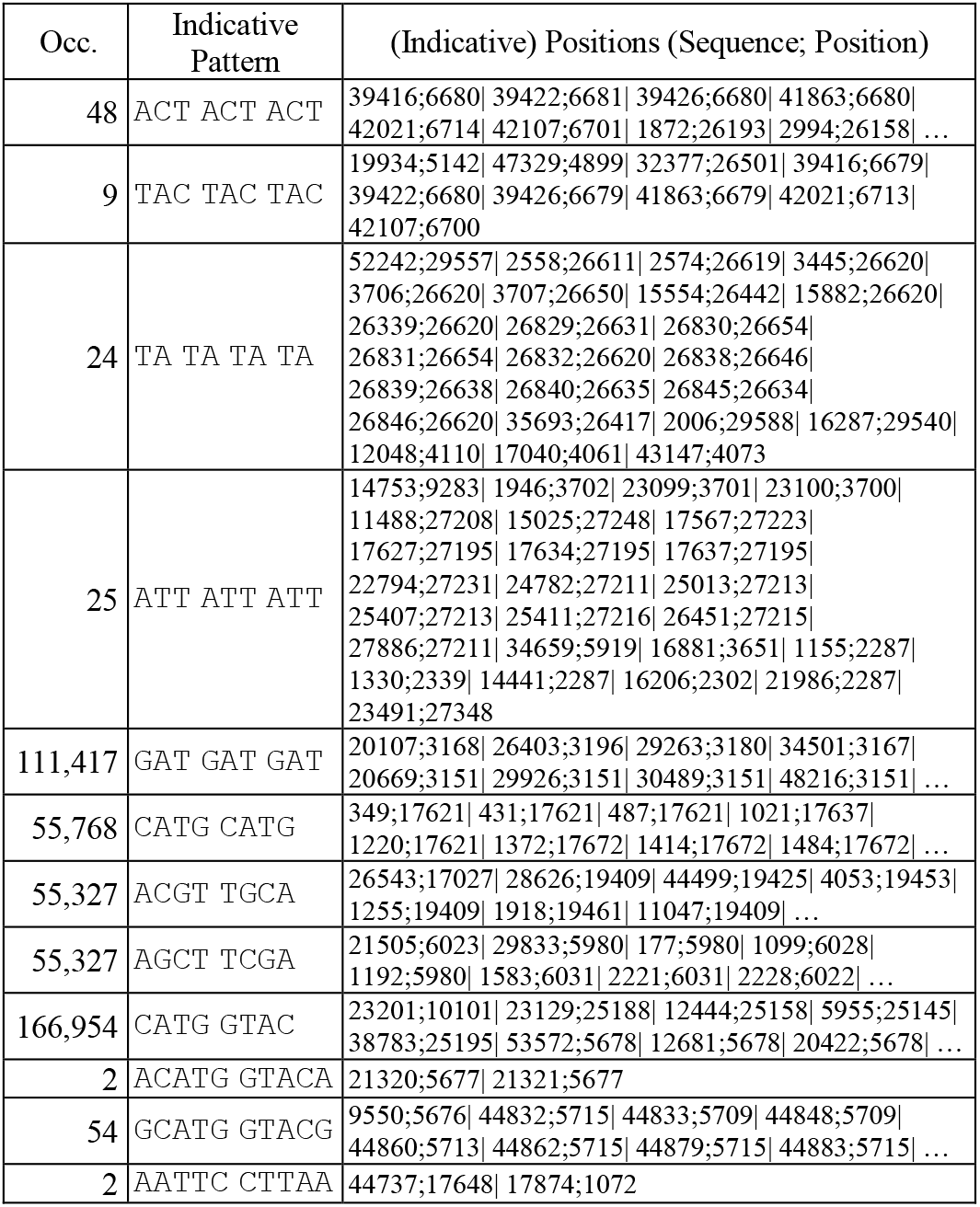
Palindromes and Tandems in SARS-CoV-2

## VI. Conclusions

The current paper presents data structures and algorithms specifically created for advanced text mining and pattern detection on discrete sequences that are adapted for biological sequences. More particularly, the purpose of the paper is to present a proof of concept and technology of the aforementioned algorithms, specifically for use on big data, with the analysis of more than 55 thousand variants of the complete SARS-CoV-2 genome. Using ordinary computers, it has been presented that it is possible to perform advanced pattern detection and produce results that can be fed as input to algorithms or used indirectly from other methodologies to perform even more detailed or diverse meta analyses. More accurately, it has been presented that with the use of LERP-RSA data structure and the single execution of ARPaD algorithm all repeated patterns can be detected, forming a vast database of results that algorithms such as SPaD and MPaD can filter and explore to perform several meta analyses. Both LERP-RSA data structure and ARPaD algorithm are very efficient and can produce the results in a few hours using commodity hardware while SPaD and MPaD can perform various analyses in few seconds.

The purpose of the current work is to unveil the potential benefits from the use of LERP-RSA and ARPaD for bioinformatics and computational biology purposes. In future work a more detailed and thorough description on particular problems will be presented with more custom-made methodologies and algorithmic variations.

